# Divergent transcriptional and transforming properties of PAX3-FOXO1 and PAX7-FOXO1 paralogs

**DOI:** 10.1101/2021.08.18.456823

**Authors:** Line Manceau, Julien Richard Albert, Pier-Luigi Lollini, Maxim V. C. Greenberg, Pascale Gilardi-Hebenstreit, Vanessa Ribes

## Abstract

The hallmarks of the alveolar subclass of rhabdomyosarcoma are chromosomal translocations that generate chimeric PAX3-FOXO1 or PAX7-FOXO1 transcription factors. Both PAX-FOXO1s result in related cell transformation in animal models, but both mutations are associated with distinct pathological manifestations in patients. To assess the mechanisms underlying these differences, we generated isogenic fibroblast lines expressing either PAX-FOXO1 paralog. Mapping of their genomic recruitment using CUT&Tag revealed that the two chimeric proteins have distinct DNA binding preferences. In addition, PAX7-FOXO1 causes stronger *de novo* transactivation of its bound regions than PAX3-FOXO1, resulting in greater transcriptomic dynamics involving genes regulating cell shape and cycle. Consistently, PAX3-FOXO1 accentuates fibroblast cellular traits associated with contractility and surface adhesion and limits entry into M phase. In contrast, PAX7-FOXO1 drives cells to adopt an amoeboid shape, reduces entry into S phase, and causes more genomic instabilities. Altogether, our results argue that the diversity of rhabdomyosarcoma manifestation arises, in part, from the divergence between the transcriptional activities of PAX3-FOXO1 and PAX7-FOXO1. Furthermore, the identified pronounced deleterious effects of PAX7-FOXO1 provide an explanation for the low frequency of the translocation generating this factor in patients with rhabdomyosarcoma.

## Introduction

Fusion-positive rhabdomyosarcomas (FP-RMS) are one of the most metastatic and deleterious subgroups of paediatric soft tissue cancers [1]. Their emergence and development are closely associated with the activity of two paralogous fusion transcription factors (TFs), PAX3-FOXO1 and PAX7-FOXO1 [1]. They are generated by the chromosomal translocations t(2;13)(q35;q14) and t(1;13)(p36;q14), which fuse the paired DNA-binding (PrD) and homeodomain (HD) domains of PAX3 or PAX7 to the transactivation domain of FOXO1. Patients with t(2;13) or t(1;13) translocations have broadly similar symptomatic features, with clusters of round cells with sparse cytoplasm separated by fibrous septa [2]. In contrast, patients with the PAX3-FOXO1 translocation are more frequent, generally older, tend to harbour more metastases, and have poorer survival rates than patients expressing PAX7-FOXO1 [3].

These clinical differences could arise from distinct and nonexclusive mechanisms. First, the specific anatomic positions of PAX3-FOXO1 and PAX7-FOXO1 tumours would argue for divergences in the cell of origin (e.g.[4]). Second, genetic aberrations that accumulate following t(2;13) or t(1;13) translocations segregate PAX3-FOXO1 and PAX7-FOXO tumours and may orient tumours towards distinctive states [5,6]. Both tumour types often undergo genome-wide duplication and carry focal chromosomal amplifications. While these amplifications extend the chromosomal segment carrying the PAX7-FOXO1 fusion, they increase the copy number of pro-proliferative genes in PAX3-FOXO1 tumours. Finally, each of the PAX-FOXO1s could have its own transcriptional activity. Supporting this idea, PAX3-FOXO1 and PAX7-FOXO1 RMS differ in their DNA methylation profiles and transcriptome [7–9].

PAX3-FOXO1 and PAX7-FOXO1 have been shown, using reporter transgenes, to possess much greater transactivation potential than the normal TFs PAX3 and PAX7 [10] and to generate, when combined with pro-proliferative mutations, RMS-like growth masses in several animal models [11–14]. The molecular mechanisms underlying their tumorigenic transfection activity were mainly studied using PAX3-FOXO1 as a model [15–18]. PAX3-FOXO1 binding to non-coding *cis*-regulatory modules (CRMs) is mediated by its PrD alone or in combination with its HD, but rarely by its HD alone [15,16]. Furthermore, PAX3-FOXO1-bound CRMs are enriched in E-box motifs recognized by bHLH TFs [15,16]. This is consistent with PAX3-FOXO1 acting with other TFs, including myogenic bHLH TFs in establishing central active CRMs in RMS cells [15,19]. In addition, the bHLH TF MYOD1 has been shown to increase the transactivation potential of PAX7-FOXO1 as measured using transcriptional reporters [20], while both PAX-FOXO1 inhibit MYOD1-mediated myogenic differentiation [21]. Finally, characterization of the chromatin landscapes of PAX3-FOXO1-bound CRMs in FP-RMS cell lines and fibroblasts revealed that PAX3-FOXO1 can bind and open previously closed chromatin loci, which acquire the characteristics of potent transactivating enhancers (super enhancers) [15]. Importantly, the mechanisms by which the two fusion TFs regulate transcription have not yet been systematically compared [7,10]. Therefore, it is not known whether the recruitment and genomic trans-activity of PAX7-FOXO1 is reminiscent of PAX3-FOXO1.

To gain insights into the role of the PAX-FOXO1 fusion TFs in tumorigenesis and heterogeneity, we directly compared their activities. By combining transcriptomic and chromatin profiling experiments with cell morphology and cell cycle assays, our results revealed a divergent transforming potential between the two PAX-FOXO1s, based on both differential use of their DNA binding domains and distinct trans-activation potential.

## Results & discussion

### Divergent genomic occupancy and transactivation potential of PAX3-FOXO1 and PAX7-FOXO1

To compare the activities of the PAX3-FOXO1 and PAX7-FOXO1 fusion proteins, we sought to design a system in which their expression can be controlled and expressed at similar levels in an identical cellular context. Human foreskin fibroblasts (HFF) were engineered to express a copy of a FLAG-tagged version of these TFs in a doxycycline (DOX)-inducible manner from the *AAVS1* safeguard locus (Figure S1b). Three independent cell lines expressing PAX3-FOXO1 or PAX7-FOXO1 were used. Their phenotype and that of control HFF cells was analysed 48 hours after DOX exposure. Under this condition, all three PAX3-FOXO1 and PAX7-FOXO1 lines expressed similar fusion protein levels in bulk and at the single cell level (Figure S1c-e).

Using these cell lines, we first mapped the genomic targets of PAX3-FOXO1 and PAX7-FOXO1 using Cleavage Under Targets and Tagmentation (CUT&Tag) experiments [22] (Figure 1; S2). We used two separate antibodies to detect the fusion TFs directed against either the N-terminal FLAG tag or the C-terminal FOXO1 domain. The latter could be used, as wild-type FOXO1 levels were low in HFFs (Figure S1c-d). Data obtained with these two antibodies and in separate cell lines gave similar results, validating the approach (Figure S2a). In total, we identified 6000 CRMs with PAX3-FOXO1 and/or PAX7-FOXO1 binding signals (Figure 1a,b; S2a; Table S1). The majority of PAX-FOXO1s bound CRMs occupied distal intronic or intergenic regions (Figure S2b) and contained DNA motifs known to be recognized by PAX3 and PAX7 DNA-binding domains (Figure 1c-i-iv, S2c; Table S2).

**Figure 1:**
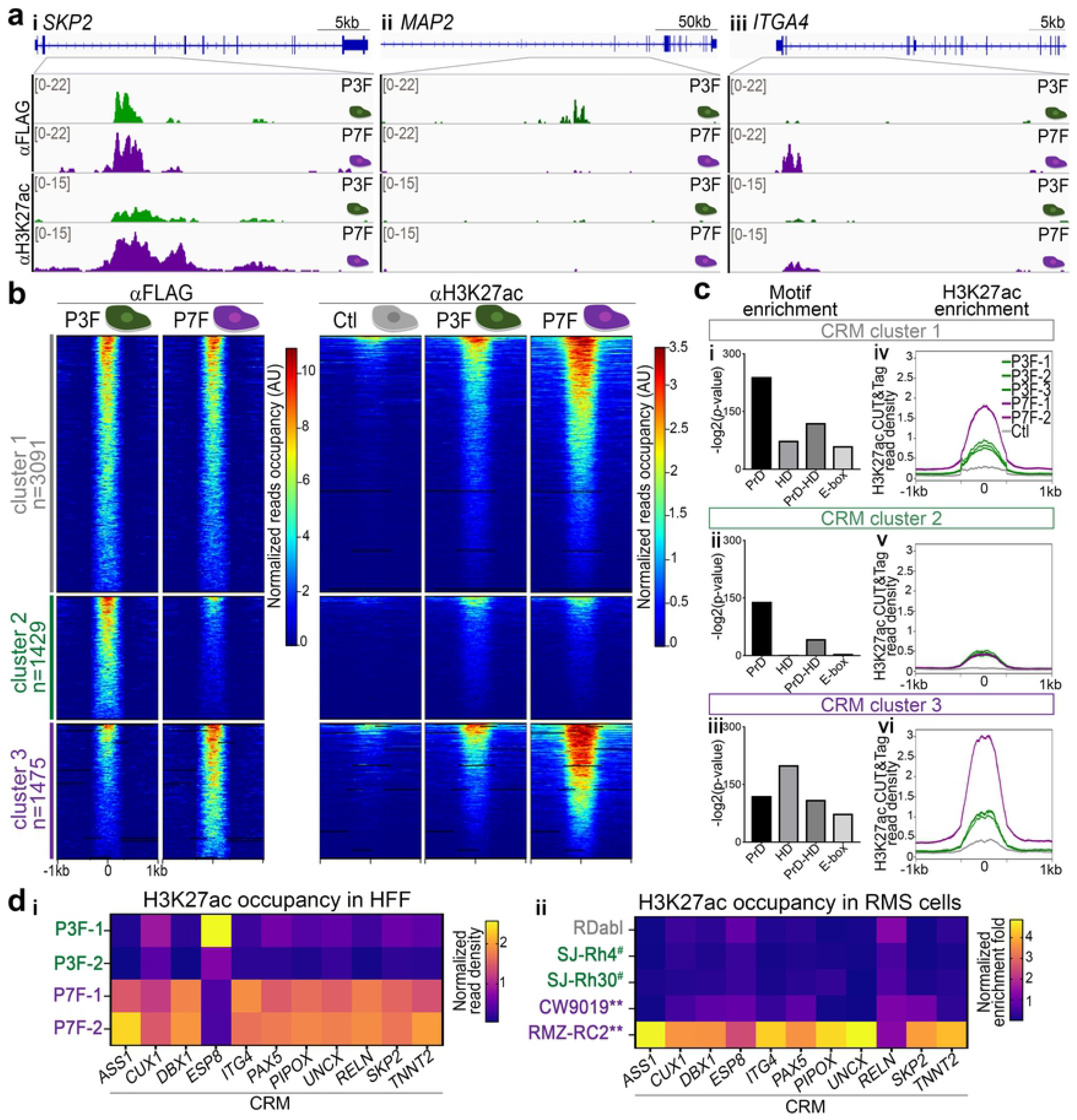
PAX3-FOXO1 and PAX7-FOXO1 recruitment to the genome and impact on H3K27ac deposition. **(a)** Examples of IGV tracks showing normalized FLAG and H3K27ac CUT&Tag reads distribution in cells expressing either PAX3-FOXO1 (P3F) or PAX7-FOXO1 (P7F) 48h post-DOX treatment. Scales in counts per million (CPM). **(b)** Heatmaps of normalized FLAG and H3K27ac CUT&Tag signals obtained in control (Ctl), PAX3-FOXO1 (P3F) and PAX7-FOXO1 (P7F) expressing HFF 48h post-DOX treatment in three CRM clusters. Cluster 1 contains CRMs onto which the occupancy rate of PAX3-FOXO1 and PAX7-FOXO1 are similar. Cluster 2 CRMs are more bound by PAX3-FOXO1 than PAX7-FOXO1, and cluster 3 is the converse of cluster 2. **(c)** Left panels: Enrichment for DNA binding motifs recognized by PAX paired (PrD) and/or homeodomain (HD) and by bHLH TFs (E-BOX) in the CRMs belonging to the clusters defined in (b) (bars: -log2(*p*-value)). Right panels: Average profiles of normalized H3K27ac CUT&Tag signals at CRMs belonging to the clusters defined in (b) in the indicated control (Ctl), P3F or P7F cell lines. Note that the two P7F purple curves are superimposed. **(d)** Heatmaps showing the relative levels of H3K27ac CUT&Tag signals on PAX-FOXO1s bound CRMs nearby the indicated genes in either HFF expressing PAX3-FOXO1 or PAX7-FOXO1 (left heatmap) or indicated RMS cell lines (right heatmap, FN-RMS lines are in grey, PAX3-FOXO1 FP-RMS in green and marked with ^#^ and PAX7-FOXO1 FP-RMS in purple and labelled with **). The left heatmap was generated using sequencing reads, while the right heatmap was established from quantitative RT-PCR data. Fold changes across samples are colour coded from purple-blue (lower levels) and yellow (higher levels).

For approximately half of these CRMs, the binding signals of PAX3-FOXO1 and PAX7-FOXO1 were comparable (Figure 1a-i, cluster 1 in 1b). In contrast, 1429 CRMs showed increased occupancy for PAX3-FOXO1 relative to PAX7-FOXO1 (Figure 1a-ii, cluster 2 in 1b) and 1475 CRMs the opposite (Figure 1a-iii, cluster 3 in 1b). Whereas PAX7-FOXO1 binding was barely detected in CRMs preferentially bound by PAX3-FOXO1, PAX3-FOXO1 recruitment was detected on most CRMs preferentially bound by PAX7-FOXO1 (Figure 1b). Hence, the two paralog proteins have divergent recruitment patterns in the genome, with PAX3-FOXO1 being more widespread than PAX7-FOXO1.

To test whether this may stem from a different use of their DNA-binding domains [23,24], we probed the different sets of CRMs for several PAX3/7- and PAX3-FOXO1-specific PrD, PrD+HD, and HD DNA motifs (see Materials and Methods; Figure S2c; Table S2). All PAX-FOXO1-bound CRMs were enriched in motifs recognized by PrD and, to a lesser extent, by PrD+HD (Figure 1c-i-iii). Motifs recognized by the HD alone were detected in CRMs bound by both PAX-FOXO1s (Figure 1c-i). However, this motif was sparsely present in PAX3-FOXO1 CRMs (Figure 1c-ii) and, in contrast, most represented in PAX7-FOXO1 CRMs (Figure 1c-iii). Overall, these results indicate that PAX3-FOXO1 uses its PrD and HD to bind DNA, whereas PAX7-FOXO1 preferentially uses its HD [23,24]. Thus, the fusion between the DNA-binding domains of PAX and the transactivation domain of FOXO1 preserved the diversification that has occurred during evolution in the modes of recruitment to the genome of PAX3 and PAX7. We then scanned PAX-FOXO1-related CRMs for E-BOX motifs recognized by class II bHLH TFs (see Materials and Methods; Figure S2c; Table S2). Apart from the PAX3-FOXO1-specific CRMs, this E-BOX was strongly represented in the other PAX-FOXO1s-bound CRMs, supporting a common interaction between bHLH TFs and both PAX-FOXO1s [15,19–21] (Figure 1c-i-iii).

We next assessed the activity status of PAX-FOXO1-related regions by mapping the distribution in the genome of the active histone mark H3K27ac using CUT&Tag (Figure 1a-c). Recruitment of PAX3-FOXO1 and PAX7-FOXO1 to the genome was generally correlated with *de novo* deposition of H3K27ac (Figure 1a,b,c,iv-vi). Therefore, both factors induce active chromatin signatures in healthy cells [15]. Intriguingly, in all CRMs, H3K27ac deposition was much more increased by PAX7-FOXO1 than by PAX3-FOXO1 (Figure 1a,b, Figure 1c-iv-vi). Even in PAX3-FOXO1-specific CRMs where PAX7-FOXO1 is poorly recruited, H3K27ac occupancy levels in PAX7-FOXO1 cells were as high as in PAX3-FOXO1 cells (Figure 1c-v). Thus, PAX7-FOXO1 exhibits a higher transactivation potential than PAX3-FOXO1. This is reinforced by a principal component analysis of H3K27ac occupancy in the 6000 PAX-FOXO1 bound CRMs showing that PAX7-FOXO1 set up a chromatin state more distant from that of control cells than PAX3-FOXO1 (Figure S2d).

To further test this idea, we compared, on a few identified CRMs, the deposition of H3K27ac in PAX-FOXO1-expressing HFFs to the recruitment of this mark in RMS cell lines (Figure 1d). For the latter, we employed CUT&Tag assays and used quantitative RT-PCR to assess the enrichment fold for H3K27ac. We used the fusion-negative RDabl cells, SJ-Rh30 and SJ-Rh4 cells expressing PAX3-FOXO1 and the CW9019 and RMZ-RC2 line expressing PAX7-FOXO1 [25]. The levels of fusion proteins in SJ-RH30, SJ-RH4 and RMZ-RC2 cells were high and comparable [14,26], whereas CW9019 had much lower levels of PAX7-FOXO1 [25]. We selected 11 CRMs following 4 criteria. They were i) bound by PAX3-FOXO1 or PAX7-FOXO1 in PAX-FOXO1 HFF lines (Table S1), ii) shown to be occupied by PAX3-FOXO1 in RMS cell lines [15], iii) nearby genes with higher expression in FP-RMS than in FN-RMS [14] and iv) close to genes with PAX-FOXO1-induced expression in HFF lines (Figure 2a, Table S3). In PAX-FOXO1 HFF lines, 10 of these 11 CRMs displayed higher levels of H3K27ac occupancy in cell lines expressing PAX7-FOXO1 than in those expressing PAX3-FOXO1 (Figure 1di). Strikingly, H3K27ac occupancy on all of these CRMs, with the exception of the *RELN* CRM, appeared at least 4-fold higher in RMZ-RC2 cells than in all other RMS cells (Figure 1dii). Furthermore, on CRMs adjacent to *DBX1, PAX5*, and *SKP2*, H3K27ac enrichment was also significantly higher in CW9019 cells weakly expressing PAX7-FOXO1 compared to PAX3-FOXO1 and RDabl cells. Thus, PAX7-FOXO1 also emerges as a more potent transactivator than PAX3-FOXO1 in fully transformed contexts. Altogether, our results demonstrate that the two RMS-associated fusion TF paralogs displayed divergent genome recruitment, likely stemming from previously identified discrete affinities of PAX3 and PAX7 DNA-binding domains for specific DNA motifs [23,24], and their differential trans-activation potential.

**Figure 2:**
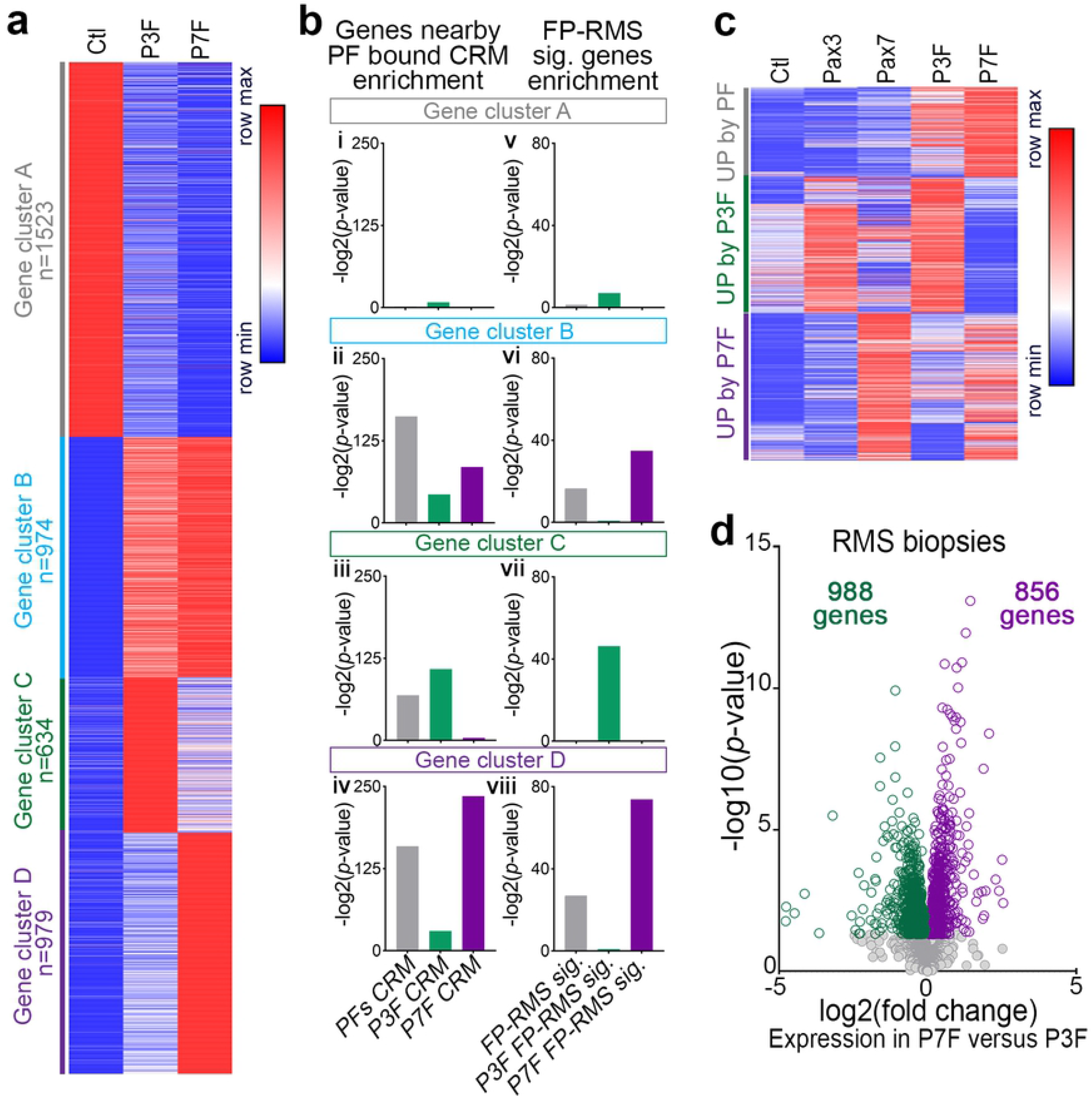
Characterization of PAX3-FOXO1 and PAX7-FOXO1 specific transcriptomic signatures. **(a)** Heatmap of 4 K-means clusters of genes showing significant variations in their expression between control, PAX3-FOXO1 (P3F) and PAX7-FOXO1 (P7F) expressing cells. Fold changes across samples are colour-coded in blue (minimum levels) to red (maximum levels). **(b)** Left panels: Enrichment for genes in the vicinity of CRMs defined in Figure 1b (PFs CRMs, P3F CRMs, P7F CRMs are respectively clusters 1,2,3 in Figure 1b) in the gene clusters defined in (a). Right panels: Enrichment for genes associated with FP-RMS^11^ (FP-RMS sig.) and genes associated with PAX3-FOXO1 (P3F FP-RMS sig.) and PAX7-FOXO1 (P7F FP-RMS sig.) in RMS biopsies in the gene clusters defined in (a). Bars: -log2(*p*-value). **(c)** Heatmap of K-means clustered genes that displayed higher expression in P3F and/or P7F transfected cells compared to control cells. Fold changes across Pax3, Pax7, P3F and P7F samples are colour coded in blue (lower level) and in red (higher level). **(d)** Volcano plot comparing gene expression levels assayed by microarray between PAX3-FOXO1 (P3F) and PAX7-FOXO1 (P7F) biopsies, with statistical significance (-log10(*p*-value)) on the y-axis versus the magnitude of change (log2(fold change in PAX7-FOXO1 samples compared to PAX3-FOXO1)) on the x-axis. Grey dots: not significantly differentially expressed genes, green dots: significantly upregulated genes in PAX3-FOXO1 samples, purple dots: significantly upregulated genes in PAX7-FOXO1 samples.

### PAX3-FOXO1 and PAX7-FOXO1 specific transcriptomic landscapes

To investigate the phenotypic consequences of the divergent activity of PAX3-FOXO1 and PAX7-FOXO1 on chromatin, we first compared the RNAseq-based transcriptome of HFFs expressing either chimeric TF or control HFFs (Figure 2a, S3a; Table S3). Mirroring the differential H3K27ac landscapes, the transcriptome of PAX7-FOXO1 cells diverged more from controls than the transcriptome of PAX3-FOXO1 cells (Figure S3a). We identified the genes whose expression varied the most between samples, which likely underlay the segregation of the transcriptome, and used K-means clustering heatmaps to highlight their behaviour in the samples (Figure 2a). PAX3-FOXO1 and PAX7-FOXO1 decreased the expression of the same subset of genes (cluster A in Figure 2a; Table S3). This subset was weakly enriched in genes in the vicinity of the identified PAX-FOXO1 recruitment sites, and was therefore unlikely to be directly regulated by PAX-FOXO1s (Figure 2bi). In contrast, the three clusters of genes upregulated by PAX3-FOXO1 and/or PAX7-FOXO1 were significantly enriched in genes nearby PAX-FOXO1s bound CRMs (clusters B to D in Figure 2a;b-ii-iv). In addition, genes more induced by PAX7-FOXO1 than by PAX3-FOXO1 were mainly enriched in CRMs preferentially bound by PAX7-FOXO1 (cluster D in Figure 2a;b-iv). Conversely, genes on which PAX3-FOXO1 had a greater impact than PAX7-FOXO1 were predominantly enriched in CRMs bound by PAX3-FOXO1 (cluster C in Figure 2a;b-iii). Thus, differences in PAX-FOXO1 recruitment are responsible for differential gene activation. We next compared the transcriptomes of cells transiently expressing murine *Pax3, Pax7*, or *PAX3-FOXO1* or *PAX7-FOXO1*. Genes induced by both PAX3-FOXO1 and PAX7-FOXO1 were poorly upregulated by Pax3 and Pax7. In contrast, genes that were specifically induced by one of the PAX-FOXO1 paralogs were also induced by its wild-type PAX variant and were not induced by the PAX paralog variant (Figure 2c, S3b). Thus, the induction of specific gene cohorts by one of the PAX-FOXO1 chimeras is a property that emanates from its PAX moiety, whereas the induction of common genes is a property of the fusion protein.

We next wondered whether gene sets specifically associated with the activity of one PAX-FOXO1 in HFF could explain some of the variation in transcriptomic status of RMS tumours [8,9,27]. We first evaluated the specific gene signatures of the PAX-FOXO1 paralog in RMS using previously published microarray data on 99 PAX3-FOXO1 and 34 PAX7-FOXO1 biopsy samples [14] (Figure 2d; Table S4). We identified 988 genes more enriched in PAX3-FOXO1 biopsies than in PAX7-FOXO1 biopsies and 856 genes specifically enriched in PAX7-FOXO1 tumours. Functional annotation of these genes highlighted that the PAX3-FOXO1-specific signature was enriched in regulators of cell cycle, cell migration, and metabolism (Figure S4a, Table S5). The genes associated with PAX7-FOXO1, on the other hand, encoded regulators of embryonic lineage differentiation as well as genes involved in cytoskeletal remodelling (Figure S4b; Table S5). Thus, paralog-specific transcriptional states could confer particular cellular traits to the cell transformation. Importantly, enrichment analyses indicated that PAX3-FOXO1 and PAX7-FOXO1 specific gene signatures in RMS and HFFs were significantly overlapping (Figure 2b-v-viii). Hence, PAX-FOXO1-dependent genetic signatures established in the cell of origin would be preserved despite the accumulation of genetic aberrations during cell transformation [5,6] and could provide specific tumorigenic features [3]. Consistent with this idea, the function of genes differentially expressed between PAX3-FOXO1- and PAX7-FOXO1-expressing HFFs was reminiscent of those differentially expressed between (t2;t13)- and (t1;t13)-carrying RMS cells (Figure S4c,d; Table S5).

### Deeper alterations of cell architecture and cell cycle induced by PAX7-FOXO1 than by PAX3-FOXO1

Genes whose expression was differentially modulated by PAX3-FOXO1 or PAX7-FOXO1 were enriched in cell cycle and cell shape regulators (Figure S4,5). As such, control, PAX3-FOXO1 and PAX7-FOXO1-expressing samples differentially expressed specific members of the Integrin, Cadherin, Semaphorin, small GTPases, Guanine nucleotide exchange factor (GEF), GTPase-activating protein (GAP) families or actin-binding and processing proteins (Figure S5a; Table S3). Hence, they likely possess their own toolkit for regulating acto-myosin network dynamics and cell adhesions (Figure S5a). Accordingly, cell morphology in these samples was distinct. This was revealed by monitoring the functional architecture of the cytoskeleton using phalloidin-labeled F-actin and focal adhesions immunostained with anti-Paxillin (Figure 3a, S6i-v). While control and PAX3-FOXO1-expressing cells predominantly exhibited a spindle or triangular shape, more than 35% of PAX7-FOXO1 cells became rounded (Figure 3ai-ix, S6i-v). Stress fibres terminated by focal adhesions were barely visible in PAX7-FOXO1 cells (Figure 3a-iv’, S6iv,v); instead, these cells displayed filopodia-like F-actin microspikes (arrowheads in Figure 3a-iv’). In contrast, in cells expressing PAX3-FOXO1, stress fibers were greater in number and thicker than in control cells (Figure 3aiii’, S6ii,iii). This indicates that each PAX-FOXO1 paralog differentially alters actin network dynamics, with PAX3-FOXO1 favouring contractile actomyosin bundles. Further demonstrating the PAX-FOXO1 paralog specific cell shape dynamics were variations in the nucleus shape, one of the key organelles of cell proprioception [28] (Figure 3av-viii, x-xi). Expression of both PAX-FOXO1 chimeric proteins is associated with the enlargement of HFF nuclei (Figure 3a-x). Yet, consistent with the loss of actin-based tension in PAX7-FOXO1 cells, their nuclei lost their roundness and adopted a bean or multilobed shape (Figure 3a-viii, xi). Overall, these results indicate that PAX3-FOXO1 enhances cellular features related to cell contractility and surface adhesions already present in HFF. Instead, PAX7-FOXO1 orients cells towards a distinctive amoeboid-like morphological state. This would certainly lead to two discrete modes of tissue invasions [29].

**Figure 3:**
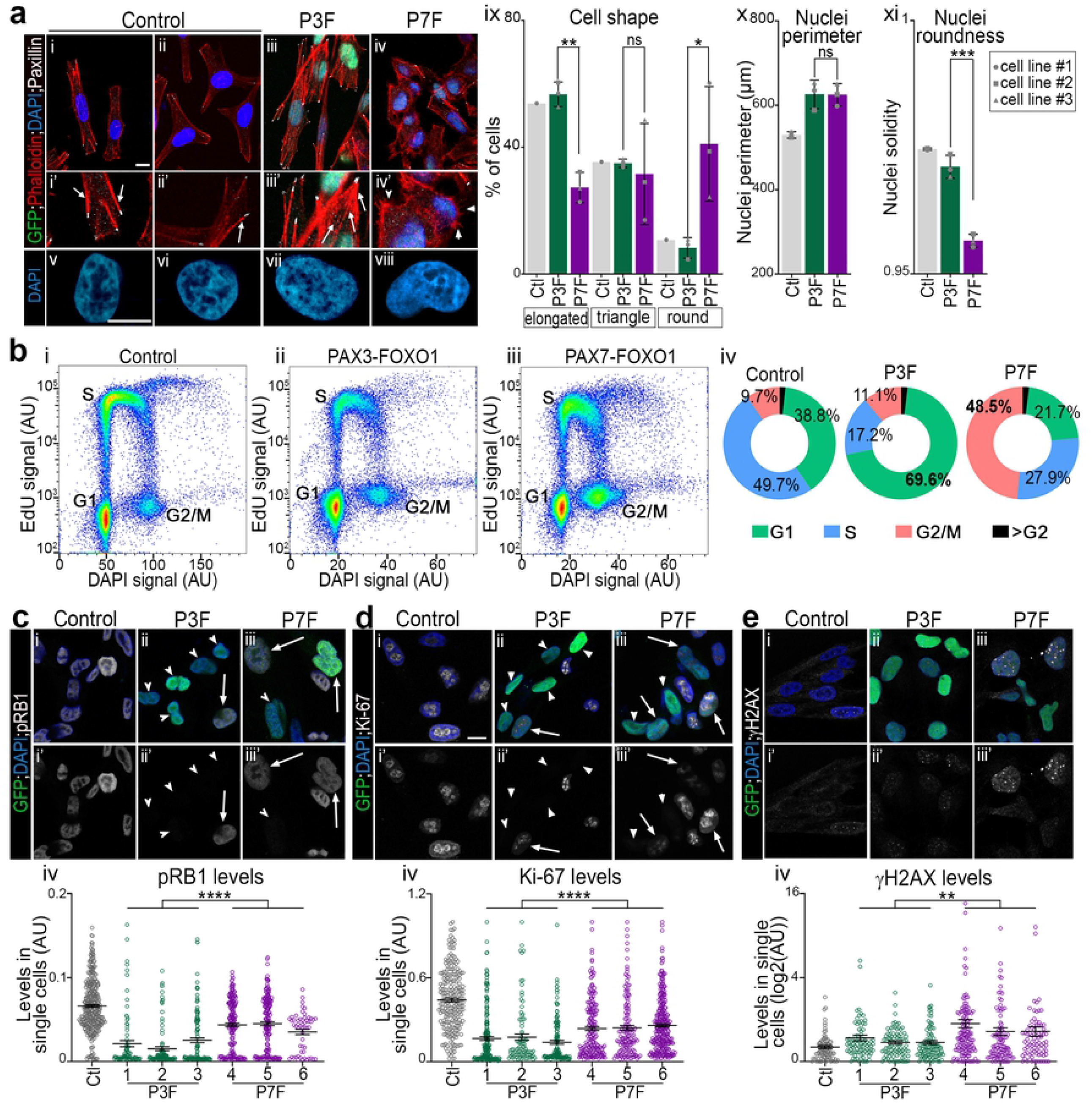
PAX3-FOXO1 & PAX7-FOXO1 activities differentially impact the shape of cells and their cycling behaviour. **(a)(i-viii)** Immunodetection of Paxillin and GFP, phalloidin based F-actin labelling and DAPI staining on the indicated HFF cell lines treated for 48h with DOX. **(ix-xi)** Proportion of cells harbouring the indicated cell morphology (ix), quantification of nuclei perimeter (x) and solidity (xi) in the indicated cell lines treated with DOX for 48h (bars: mean ± s.d.; dots: mean value in independent cell lines; Mann-Whitney U test p-value: *: *p*< 0.05, **: *p*< 0.01, ***: *p*< 0.001, ns: *p*> 0.05). **(b)(i-iii)** FACS plots displaying EdU levels and DNA content (DAPI levels) in control cells and in the GFP^+^ population of a PAX3-FOXO1 and PAX7-FOXO1 cell line treated for 48h with DOX. **(ii)** Percentage of cells in the indicated cell cycle phase established from plots in i (mean over three experiments and three independent lines). **(c-e)(i-iii’)** Immunodetection of pRB1, Ki-67, γH2AX, 53BP1 and GFP and DAPI staining on the indicated HFF cell lines treated for 48h with DOX. **(c-e)(iv)** Levels of expression of pRB1, Ki-67 and γH2AX in control, GFP^+^ PAX3-FOXO1 and GFP^+^ PAX7-FOXO1 cells treated for 48h with DOX (dots: cell values, bars: mean ± s.e.m., two-way-ANOVA *p*-values evaluating the similarities between PAX3-FOXO1 and PAX7-FOX1 cells lines: ****: *p*<0.0001).

To assess the impact of PAX-FOXO1s on the cell cycle, we labelled replicating cells with a 1h30 pulse of EdU, a thymidine analogue, prior to harvest. We then used flow cytometry to quantify DNA content and replication using DAPI and EdU levels (Figure 3b). Both PAX-FOXO1s altered the distribution of cells across cell cycle phases (Figure 3b). The number of cells in the replication (S) phase was reduced in the presence of both chimeras, but this reduction was much greater in the presence of PAX3-FOXO1 than PAX7-FOXO1 (Figure 3b). Consistently, CYCLIN-CDK-dependent phosphorylation levels of Rb1 and the cell proliferation marker Ki-67 (MKI67) were further reduced in PAX3-FOXO1 cells compared to PAX7-FOXO1 cells (Figure 3c,d, S6vi-xv). This is also in agreement with the decreased expression of core cell cycle genes and the upregulation of cell cycle inhibitors by both PAX-FOXO1 (Figure S5b). Interestingly, whereas the G1 cell population was dominant in PAX3-FOXO1-expressing HFFs, PAX7-FOXO1 expression was associated with an increased percentage of cells in G2 (Figure 3bii-iv). This observation was reinforced by the enrichment of PAX7-FOXO1 molecular signature for genes encoding regulators of the G2 to M transition, whereas PAX3-FOXO1 specifically induced inhibitors of the G1 to S transition (Figure S5b; Table S3). These distinct effects of PAX-FOXO1 paralogs on the cell cycle could underlie the frequent association in PAX3-FOXO1 tumours, but not PAX7-FOXO1, with genetic aberrations increasing the expression of MYCN and CDK4 [6], two G1-to-S transition regulators.

Finally, we tested whether PAX-FOXOs-mediated cell cycle deregulation was linked to DNA damage that can block cell cycle progression [30]. Immuno-labelling of 53BP1^+^ and γH2AX^+^ DNA double-strand break foci showed that PAX7-FOXO1 was more prone to cause genomic instabilities than PAX3-FOXO1 (Figure 3e, S6xvi-xx). It remains to be determined whether this DNA damage is mechanistically related to the enhanced transcriptional properties of PAX-FOXO1 paralogs [31] or whether it arises from the hijacking of wild-type PAX functions in preserving genomic integrity [32,33]. Yet, we hypothesize that the DNA damage and cell cycle defects associated with PAX-FOXO1 paralogs may in turn trigger the gross genomic rearrangement catastrophes that appear in the FP-RMS after PAX3-FOXO1 and PAX7-FOXO1 generating translocations [5,6,34]. In particular, the enhanced propensity of PAX7-FOXO1 to alter cell transcriptomic status, morphology, cycling, and genome stability is likely to be detrimental to cells over the long term. This could therefore explain both the silencing of PAX7-FOXO1 in most cells in RMS biopsies [35] and the lower frequency of occurrence and aggressiveness of PAX7-FOXO1-associated RMS compared with PAX3-FOXO1-associated RMS [1,3].

Overall, our study demonstrates that the two PAX-FOXO1 paralogs confer distinct molecular and cellular characteristics to healthy cells in the early stage of tumorigenic transformation, which may lead to differential manifestation of patients carrying PAX3-FOXO1 or PAX7-FOXO1 generating translocations.

## Material and methods

Due to the word limit, this section is expanded in the supplementary information file.

## Acknowledgements

We deeply thank the ImagoSeine core facility of Institut Jacques Monod, a member of France-BioImaging (ANR-10-INBS-04) and certified IBiSA. Notably, S. Many and N. Valentin for performing flow cytometry analyses and N. Moisan for training us on confocal imaging. We are grateful to L. Vinel for helping us during her undergraduate internship and V. Doye and S. Nedelec for critical inputs on our manuscript. We are thankful to people who have provided us with useful tools. We received *pAAVS1-PDi-CRISPRn* from B. Conklin and gRNAs from G. Church; ERMS and FP-RMS cell lines from C. Gauthier-Rouvière and F. Barr and HFF from M-C. Geoffroy; the anti-53BP1 and anti-γH2AX antibodies from C. Boumendil and the anti-PAX3/7HD from N. Patel. We thank Bérengère Guichard for the preparation and purification of pA-Tn5 enzyme.

## Author contributions

Conceptualization, LM, PGH, VR; Methodology, LM, JRA, PGH, VR; Software, JRA; Validation, LM, PGH, VR; Formal Analysis, LM, PGH, VR; Investigation, LM, JRA PGH, VR; Resources, JRA, PLL, MVCG, VR; Writing-Original Draft, LM, VR; Writing - Reviewing & Editing, LM, JRA, MVCG, PGH, VR; Visualization, LM, JRA, VR; Supervision, JRA, PGH, VR; Project Administration, VR; Funding Acquisition, MVCG, VR.

